# Comparing the effect of multi gradient echo and multi band fMRI during a semantic task

**DOI:** 10.1101/2024.03.20.585909

**Authors:** Ajay D. Halai, Richard N. Henson, Paola Finoia, Marta M. Correia

## Abstract

The Blood Oxygenation Level Dependent (BOLD) signal, as measured using functional magnetic resonance imaging (fMRI), is known to vary in sensitivity across the brain due to magnetic susceptibility artefacts. In particular, the ventral anterior temporal lobes (vATL) have been implicated with semantic cognition using convergent methods (i.e., neuropsychology, PET, MEG, brain stimulation) but less so with fMRI using conventional gradient-echo protocols. There are methods to alleviate signal loss but multi-echo fMRI has gained popularity. Here, additional volumes are collected that span across a range of T2* values, however, this results sub-optimum parameters (i.e., repetition times, resolution, acceleration). “Multi-band” imaging has been used with multi-echo to speed up data acquisition; however, it is unclear how these modifications contribute to fMRI sensitivity across the brain and for univariate/multivariate analyses. In the current study, we used a factorial design where we manipulated the echo and/or band to assess how well the semantic network can be detected. When comparing the precision with which activations were detected (i.e, average T-statistics), we found that multi-band protocols were beneficial, with no evidence of signal leakage artefacts. When comparing the magnitude of activations, multi-echo protocols increased activations in regions prone to susceptibility artefacts (specifically the anterior temporal lobes, ATLs). Both multi-banding and independent component analysis (ICA)-denoising of multi-echo data tended to improve multi-voxel decoding of conditions. However, multi-echo protocols reduced activation magnitude in more central regions, such as the medial temporal lobes, possibly due to higher in-plane acceleration required to collect multiple-echoes. Nonetheless, the multi-echo multi-band protocol is a promising default option for fMRI on most regions, particularly those that suffer from susceptibility artefacts, as well as offering the potential to apply advanced post-processing methods to take advantage of the increased temporal (or spatial) resolution of multi-band protocols and more principled ICA-denoising based on TE-dependence of BOLD signals.

## 1. Introduction

Gradient-echo functional magnetic resonance imaging (fMRI) using the Blood Oxygenation Level Dependent (BOLD) signal has become the dominant non-invasive tool for studying how the brain operates; however, its ability to detect signal across the whole brain is not homogenous. This can lead studies to be blind to activation within the ventral anterior temporal lobes (ATL) and orbito-frontal cortices. Bilateral ATLs have been implicated with semantic cognition across multiple convergent methods (i.e., neuropsychology, PET, MEG, brain stimulation) but less so with fMRI (see Lambon Ralph, Jefferies, Patterson, & Rogers, 2017 for a review). There have been multiple attempts to overcome these issues with fMRI, and recently multi-echo fMRI has gained popularity (see Kundu et al., 2017 for a review). Typical fMRI studies collect a single echo, which results in sensitivity to a narrow range of T2* values (ideally for maximal sensitivity to BOLD contrast); however, T2* is known to vary across the brain and between participants (Hagberg, Indovina, Sanes, & Posse, 2002). Therefore, taking multiple echoes can improve sensitivity to a range of T2* values, which is particularly important for areas with susceptibility artefacts, where signal can be detected before it dephases. There have been a number of studies comparing multi-echo with typical or modified fMRI approaches, which have reported evidence in favour of multi-echo approaches. However, there are a number of caveats associated with this literature, as discussed below.

Firstly, many studies use only a multi-echo protocol, and compare multi-echo results with those from analysing just one of the TEs derived from that protocol (usually the TE optimal for BOLD, e.g., 25-35ms for 3T; Amemiya, Yamashita, Takao, & Abe, 2019; Bhavsar, Zvyagintsev, & Mathiak, 2014; Caballero-Gaudes, Moia, Panwar, Bandettini, & Gonzalez-Castillo, 2019; Cohen, Nencka, Marc Lebel, & Wang, 2017; Cohen, Nencka, & Wang, 2018; Cohen & Wang, 2019; Dipasquale et al., 2017; Evans, Kundu, Horovitz, & Bandettini, 2015; Fernandez, Leuchs, Sämann, Czisch, & Spoormaker, 2017; Gilmore, Agron, González-Araya, Gotts, & Martin, 2022; Halai, Parkes, & Welbourne, 2015; Halai, Welbourne, Embleton, & Parkes, 2014; Heunis et al., 2021; Kovářová, Gajdoš, Rektor, & Mikl, 2022). However, while the single-echo timeseries extracted from a multi-echo sequence resembles a ‘typical’ fMRI dataset, it is likely to be inferior in quality to data from an optimised single-echo sequence. This is because multi-echo acquisition requires sequence parameters that are not necessarily optimal for a single-echo data (e.g., higher in-plane acceleration, reduced k-space sampling, etc.), which can result in aliasing or noise enhancement (e.g., Deshmane, Gulani, Griswold, & Seiberlich, 2012). Thus while comparing results from a single versus multiple echoes from the same multi-echo sequence provides tight control of other sequence parameters, thereby isolating the advantage of multi-echoes, it is not a fair comparison for practical decisions about whether to use a multi-echo versus standard single-echo protocol. Critically, we found only two studies that compared a multi-echo protocol against a typical single-echo protocol (Poser, Versluis, Hoogduin, & Norris, 2006; Kirilina, Lutti, Poser, Blankenburg, & Weiskopf, 2016); remaining studies used a multi-band accelerated single-echo protocol (Cohen, Chang, & Wang, 2021; Cohen, Jagra, Visser, et al., 2021; Cohen, Jagra, Yang, et al., 2021; Cohen, Yang, Fernandez, Banerjee, & Wang, 2021; Fazal et al., 2023; Lynch et al., 2020) and/or compared different denoising strategies (Lombardo et al., 2016). Using resting-state fMRI (rsfMRI), Poser et al. (2006) demonstrated that multi-echo data had better functional contrast-to-noise ratio (CNR) across the brain than a typical protocol, particularly in susceptibility-prone regions. In a follow up study, Kirilina et al., (2016) showed greater task activation during emotional and reward-based learning in orbitofrontal cortices, most likely due to effective recovery of signal in these susceptible areas, but less activation in deep brain regions, most likely due to higher acceleration and/or smaller voxel size. Similarly, Fazal et al., (2023) showed that multi-echo multi-band had greater sensitivity in areas affected by signal inhomogeneity but multi-band only did show greater sensitivity in visual areas (not deep brain regions).

Secondly, another recent advance in echo planar imaging (EPI), often referred to as simultaneous multi-slice (SMS) or “multi-band” imaging, has made it possible to acquire whole-brain fMRI datasets with spatial and/or temporal resolution that are several times higher than typical protocols (Moeller et al., 2010; Setsompop et al., 2012). Multi-band imaging has been shown to reduce temporal aliasing of high frequency noise sources, increase statistical power, and improve resting-state network estimation/reliability (Feinberg et al., 2010; Griffanti et al., 2014; Smith, Beckmann, Andersson, Auerbach, Bijsterbosch, Douaud, Duff, Feinberg, Griffanti, Harms, Kelly, Laumann, Miller, Moeller, Petersen, Power, Salimi-Khorshidi, Snyder, Vu, Woolrich, Xu, Yacoub, Uǧurbil, et al., 2013). Studies have also shown that higher temporal resolution leads to a greater number of independent components identified in resting state data, and this benefit is lost when down-sampling the multiband series to match a typical protocol (Olafsson, Kundu, Wong, Bandettini, & Liu, 2015). However, fMRI sequences with very high multi-banding acceleration can also impair SNR (e.g., Chen et al., 2015; Smith et al., 2013; Demetriou et al., 2018). Moreover, while multi-band modifications have mainly been promoted for improved estimation of resting-state connectivity (Smith, Beckmann, Andersson, Auerbach, Bijsterbosch, Douaud, Duff, Feinberg, Griffanti, Harms, Kelly, Laumann, Miller, Moeller, Petersen, Power, Salimi-Khorshidi, Snyder, Vu, Woolrich, Xu, Yacoub, Uğurbil, et al., 2013; Uğurbil et al., 2013), their benefit for task-based fMRI analysis has been less clear (Demetriou et al., 2018; Todd et al., 2017), since low-frequency fMRI noise is unlikely to be correlated (phase-locked) with the task regressors, and so can be removed by high-pass filtering the data. Moreover, a potential disadvantage of multi-band acquisition is false-positives due to signal leakage across simultaneously excited slices (Xu et al., 2013). Todd et al. (2016) found that slice leakage was particularly apparent during multi-band factors greater than 4. Despite these caveats, studies have combined multi-band with multi-echo imaging, to compensate for the decreased spatial and/or temporal resolution resulting from collecting additional TEs. This may partially explain why most studies focus on comparing a multi-band, multi-echo protocol to a multi-band, single-echo protocol. However, it is difficult to disentangle the effects of echo and band without both elements being manipulated independently.

Thirdly, the criteria used to determine whether multi-echo protocols add benefit varies in the literature. Many studies use rs-fMRI to estimate signal quality through measures like temporal signal-to-noise ratio (tSNR) or CNR, although some studies have investigated changes to functional connectivity/networks. In contrast, task-based fMRI studies have focused on statistical results in primary sensory cortices (using, e.g., visual checkerboard, finger tapping, breath hold). However, when comparing fMRI protocols, it is important to employ tasks that are known to produce reliable activations across many brain regions, in order to span regions with both high and low levels of susceptibility artefacts. Semantic cognition is one such function, because it activates a large language-related network across bilateral temporal and frontal lobes, in particular the anterior lateral temporal lobe (ATL; see Lambon Ralph et al., 2017 for a review). Both PET (Devlin et al., 2000) and distortion-corrected, spin-echo fMRI studies (e.g., Binney, Embleton, Jefferies, Parker, & Lambon Ralph, 2010; Humphreys, Hoffman, Visser, Binney, & Lambon Ralph, 2015; Visser, Embleton, Jefferies, Parker, & Ralph, 2010) have shown ATL activation during semantic cognition tasks that is harder to detect with gradient-echo fMRI, owing to high levels of susceptibility artefact in this region.

Here we examined the effects of protocol both on activation magnitude (e.g., percent signal change difference between two task conditions) and activation “precision”, which is a noise-normalised measure, i.e. difference in activation magnitude divided by an estimate of uncertainty in the estimate of that difference (equivalent to a T-statistic). A priori, we would expect ME protocols to recover signal in regions prone to susceptibility artefacts, which should be apparent in greater effects of task on activation magnitude (which may also translate into higher activation precision too, if noise is unaffected by the protocol). MB protocols, on the other hand, should improve activation precision by virtue of a greater number of volumes (data points) and reduced aliasing of high-frequency noise, but should not affect magnitude (since the signal magnitude is independent of how often it is sampled). We tested the reliability of any differences between protocols in terms of T-tests across participants on both activation magnitude and activation precision estimates within each participant. Note that group-level statistics are normally performed on activation magnitudes, but sometimes individual-level statistics (activation precisions) are also important to optimise, e.g., in single case-studies.

Finally, all studies to date have focused on univariate activations for task-based paradigms. This approach is the most widely-used method for identifying brain regions activated during a cognitive process, in terms of differences between conditions at each voxel, or after averaging over voxels within a region. In contrast, multivariate methods (such as multivoxel pattern analysis [MVPA]) utilise patterns of activation across many voxels (often irrespective of their mean level of activation) to determine if those patterns differ between conditions (e.g., Coutanche, 2013; Davis & Poldrack, 2013). Here we estimated the ability of multi-voxel patterns to distinguish (“decode”) two conditions. There is a growing literature suggesting that MVPA can be more powerful than univariate analysis because it exploits variance and covariance between voxels, which is discarded in univariate analyses, and can allow for participant-specific differences in the patterns by analysing derived measures across participants, such as decoding ability (Davis et al., 2014).

In this study, we evaluate the performance of different fMRI protocols to detect both univariate activation and multivariate decoding during a semantic task. The study is unique in four important aspects. First, the “baseline” comparison comes from an independent run using a typical protocol that is optimised for single-band, single-echo fMRI. Second, we manipulate number of echoes and bands independently, resulting in a 2 x 2 factorial design to better separate the effects of multi-echoes and multi-banding (and any potential interaction between these factors). Third, our semantic judgement task is known to activate a network spanning areas that typically have both good and poor image quality. Fourth, we investigate the effects of fMRI protocol on activation magnitude, activation precision and MVPA. Finally, we also investigated whether multiple bands resulted in false-positives due to slice leakage, for both univariate and multivariate analyses.

## 2. Materials and Methods

### 2.1. Participants

We recruited 21 healthy native English speakers (13 females, mean age = 28.9 +-8.75 years, range 19 to 50 years). All participants were right-handed, had normal or corrected to normal vision and no known neurological disorders (i.e., dyslexia, neurodegeneration, etc). The experiment was approved by the Cambridge Psychology Research Ethics Committee (CPREC).

### 2.2. Stimuli design

Participants completed a semantic “triad” task, variants of which have previously been shown to produce robust activation the ventral anterior temporal lobes using dual-echo fMRI (Jung, Williams, Sanaei Nezhad, & Lambon Ralph, 2017) and spin-echo fMRI (Humphreys et al., 2015). In each trial, three pictures are presented for a matching task (Figure 1). In the semantic condition, the participant is required to press a left or right button to indicate which of the two pictures on the bottom of the screen has a semantic relationship with the (probe) picture on the top. For example, if the probe is stool and the options are cow and chicken, one would select the cow. In the control condition, the pictures are scrambled, and the task is a perceptual rather than semantic match, i.e., to indicate which bottom image is identical to the top image. The pictures were extracted from the Pyramid and Palm Trees test (Howard & Patterson, 1992) and Camel and Cactus test (Bozeat, Lambon Ralph, Patterson, Garrard, & Hodges, 2000). EPRIME software was used to display the stimuli and record responses.

**Figure 1.**
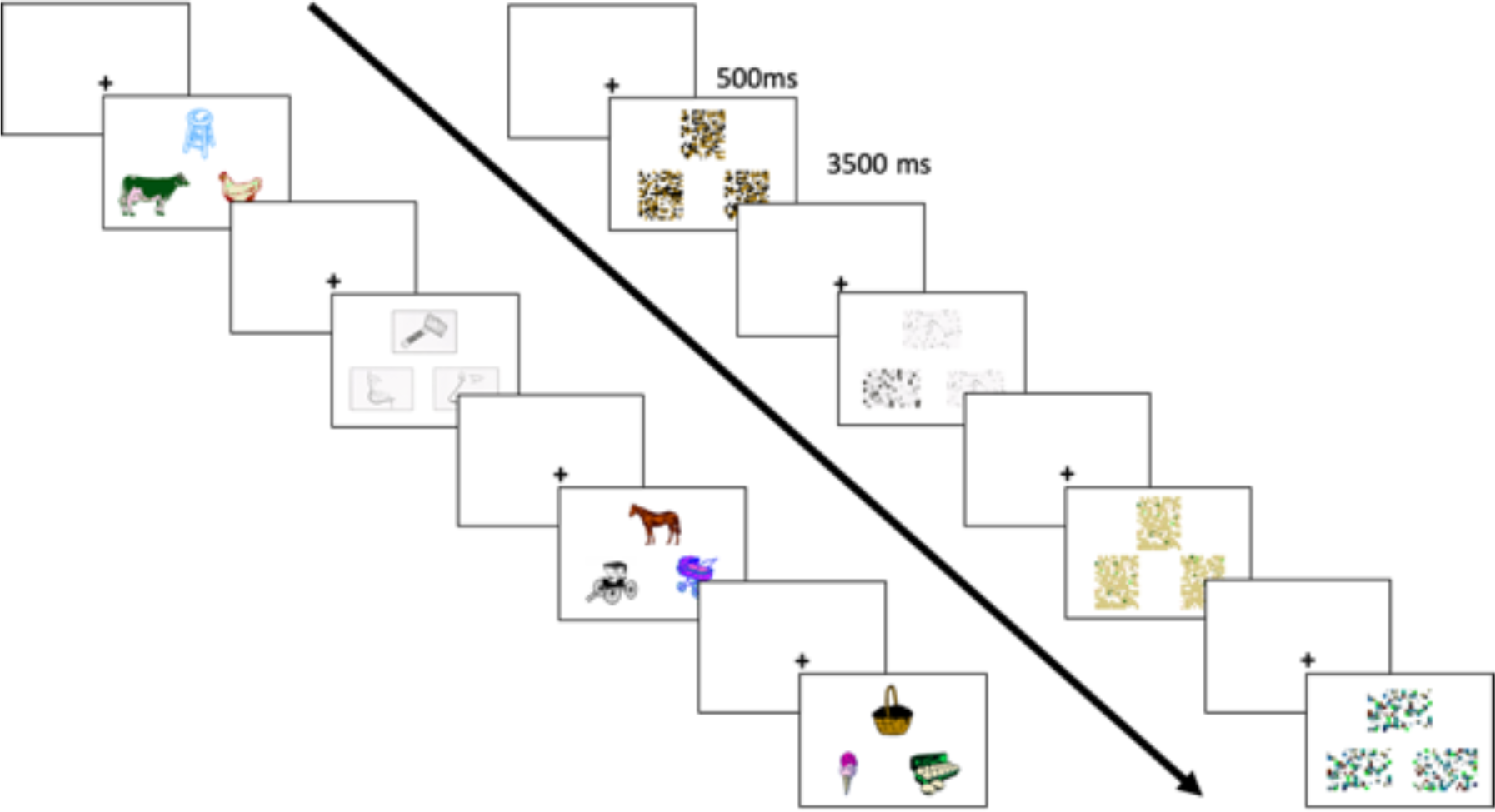
Semantic (left) and control (right) blocks of the triad task. Each block lasted 16 s that included four trials, where each trial had a 500 ms fixation and 3500 ms stimulus presentation. Participant were given a probe at the top and were asked to press the left/right button to indicate which of the two pictures below matched the probe.

We used a block design consisting of 12 replications of 3 types of blocks: semantic (S), control (C) and rest (R). The task started after a 16 s delay after the MR protocol. The 36 blocks were ordered S-C-R-C-S-R (x6) (except for the first two participants for whom the order was S-C-R x12). Each block contained four trials with the following structure: fixation for 500 ms followed by a triad for 3500 ms, resulting in a total block length of 16 s. The paradigm lasted 592 s in total including rest and was repeated for each fMRI protocol. Accuracy and reaction time were measured for each trial.

We performed a 2x2x2 ANOVA (condition x multiband x multi-echo) for accuracy and reaction time separately to test for differences in behavioural performance.

### 2.3. Data acquisition

Data were acquired using a 3T Siemens Prisma FIT scanner with a 32-channel RF head coil. The gradient-echo planar imaging (EPI) field of view (FOV) was placed approximately 30° degrees from the AC-PC line (angled away from the eyes) and full coverage of the temporal lobes was ensured by lowering the FOV at a cost of missing the top of the brain (i.e., superior parietal lobe).

We matched the four EPI protocols as much as possible across multiple parameters, while manipulating echoes and/or banding. For all EPI protocols, 80 x 80 pixel slices of 3mm x 3mm were acquired in a descending, sequential order, with a 0.45mm gap between slices (i.e., final voxel size 3 x 3 x 3.45 mm^3^). A partial Fourier = 7/8^th^ acquisition was used.

The parameters that then differed across protocols are shown in Table 1. For the MB protocols, we used a MB factor of 2, and matched spatial parameters (i.e., same slice thickness), resulting in better temporal resolution, i.e., halved TR. The flip angle was optimised for each TR. For the ME protocols, we acquired three echoes (13.00ms, 25.85ms and 38.70ms), and used a GRAPPA acceleration of 3, resulting in a slightly faster acquisition, so we increased the number of slices from 28 to 30 in order to better match the TR.

**Table 1.**
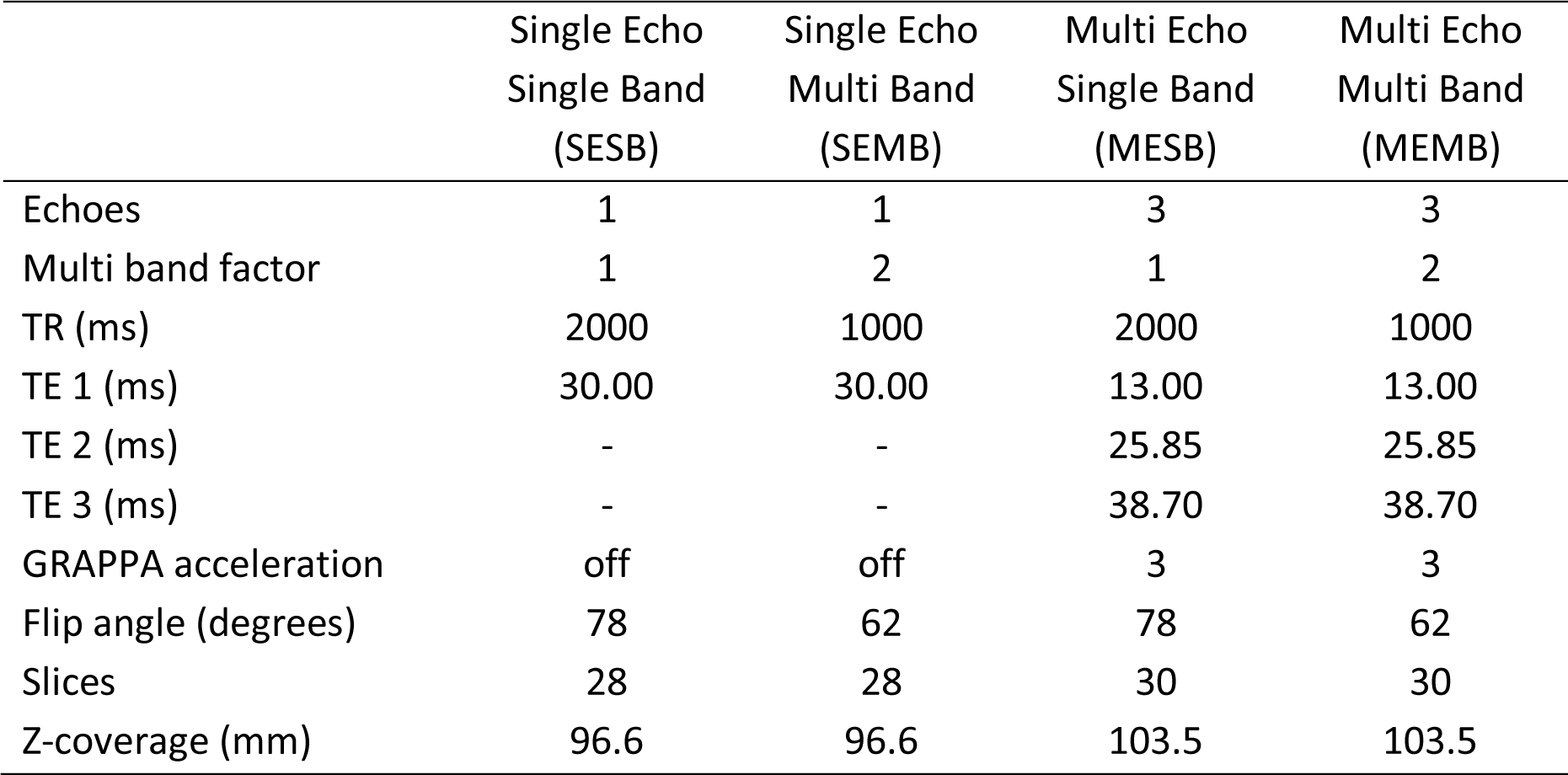
Parameters for the four EPI protocols compared in a 2x2 factorial design. Abbreviations: repetition time (TR); echo time (TE)

An MPRAGE structural image was acquired with the following parameters: TR = 2250 ms, TE = 3.02 ms, TI = 900 ms, GRAPPA = 2, FOV = 256 mm*256 mm*192 mm, voxel size = 1mm^3^, flip angle (FA) = 9°.

### 2.4. Data analysis

Data will be made publicly available upon peer review and acceptance. Code is publicly available at https://github.com/AjayHalai/Sensitivity_of_MEMB.

#### 2.4.1. Preprocessing

For reproducibility and data sharing, we converted all DICOMs to BIDS format (K. J. Gorgolewski et al., 2016) using the *HeuDiConv* tool (v0.9.0; Halchenko et al., 2020) and used an fMRIprep (v22.0.0) (Esteban et al., 2019; K. Gorgolewski et al., 2011) singularity container to process all imaging data. The only exception was for the multi-echo datasets, which were re-processed using the *tedana* tool (v0.0.11) (DuPre et al., 2021) (details below).

The T1-weighted (T1w) image was corrected for intensity non-uniformity with *N4BiasFieldCorrection* (Tustison et al., 2010), distributed with *ANTs* (v2.3.3) (Avants, Epstein, Grossman, & Gee, 2008), and used as T1w-reference throughout the workflow. The T1w-reference was then skull-stripped with a Nipype implementation of the *antsBrainExtraction.sh* workflow (from *ANTs*), using OASIS30ANTs as target template. Brain tissue segmentation of cerebrospinal fluid (CSF), white-matter (WM) and grey-matter (GM) was performed on the brain-extracted T1w using *fast* (FSL v6.0.5.1, Zhang, Brady, & Smith, 2001). Volume-based spatial normalization to standard space (TemplateFlow ID: MNI152NLin2009cAsym; Fonov, Evans, McKinstry, Almli, & Collins, 2009) was performed through nonlinear registration with *antsRegistration* (from *ANTs)*, using brain-extracted versions of both T1w and the T1w template.

For each of the functional runs, the following pre-processing was performed. First, a BOLD reference volume (from the shortest echo) and its skull-stripped version were generated using a custom methodology in fMRIPrep. Head-motion parameters with respect to the BOLD reference (transformation matrices, and six corresponding rotation and translation parameters) are estimated before any spatiotemporal filtering using *mcflirt* (Jenkinson et al. 2002). BOLD runs were slice-time corrected to the middle slice using *3dTshift* from *AFNI* (Cox & Hyde, 1997). The slice-time corrected data were corrected for head-motion (referred to as preprocessed BOLD). The BOLD reference was then co-registered to the T1w reference using mri_coreg (FreeSurfer) followed by *flirt* (Jenkinson & Smith, 2001) with the boundary-based registration (Greve & Fischl, 2009) cost-function with six degrees of freedom. Several additional confounding time-series were calculated (i.e., framewise displacement (FD), DVARS and three region-wise global signals, anatomical/temporal component-based noise correction) but we only used the six motion (translation and rotation) parameters in this study, therefore the other metrics are not described further.

For multi-echo data, we used *tedana* to reprocess the minimally pre-processed BOLD data (slice time and motion corrected) from the fMRIprep pipeline in order to obtain the ICA-denoised timeseries. A whole brain mask derived from the T1 was used to define brain space, to which a two-stage adaptive masking procedure was implemented. First, a liberal mask (including voxels with good data in at least the first echo) was used for optimal combination, T2*/S0 estimation, and denoising, while a more conservative mask (restricted to voxels with good data in at least the first three echoes) was used for the component classification procedure. A mono-exponential model was fit to the data at each voxel using nonlinear model fitting in order to estimate T2* and S0 maps, using T2*/S0 estimates from a log-linear fit as initial values. For each voxel, the value from the adaptive mask was used to determine which echoes would be used to estimate T2* and S0. In cases of model fit failure, T2*/S0 estimates from the log-linear fit were retained instead. Multi-echo data were then optimally combined using the T2* combination method (Posse et al., 1999). Principal component analysis based on the PCA component estimation with a Moving Average (stationary Gaussian) process (Li, Adalı, & Calhoun, 2007) was applied to the optimally combined data for dimensionality reduction. Next, an independent component analysis (ICA) was then used to decompose the dimensionally reduced dataset from the PCA. Kappa (kappa) and Rho (rho) values were calculated as measures of TE-dependence and TE-independence, respectively for both PCA and ICA reduced datasets. Finally, component selection was performed to identify BOLD (TE-dependent), non-BOLD (TE-independent), and uncertain (low-variance) components using the Kundu decision tree (v2.5; Kundu et al., 2013). This workflow used numpy (Walt, Colbert, & Varoquaux, 2011), scipy (Jones, Oliphant, & Peterson, 2001), pandas (McKinney, 2010), scikit-learn (Pedregosa et al., 2011), nilearn, and nibabel (Brett et al., 2019). This workflow also used the Dice similarity index (Dice, 1945; Sørensen, 1948). We retained two final timeseries from this pipeline: optimally combined T2* (t2star); and ICA-denoised (ICA-denoised).

The BOLD time-series from all protocols were resampled into standard space, generating a pre-processed BOLD run in MNI152NLin2009cAsym space. All resamplings can be performed with a single interpolation step by composing all the pertinent transformations (i.e. head-motion transform matrices and co-registrations to anatomical and output spaces). Volumetric resampling were performed using *antsApplyTransforms* (*ANTs*), configured with Lanczos interpolation to minimize the smoothing effects of other kernels (Lanczos, 1964). All images in MNI space were smoothed using 8 mm^3^ kernel in SPM12.

#### 2.4.2. 1^st^ level (within-participant) GLM

Statistical analysis was carried out using used SPM12 (v) in MATLAB r2019a to create 1^st^ level general linear models (GLM) for each protocol of interest. To be explicit, we had a 2x2 factorial design (SESB, SEMB, MESB, MEMB) but we also investigated specific effects for 1) ICA-denoised data (with “dn” suffix) and 2) a reduced multi-band dataset by extracting only odd volumes to match the number of time points in the non-accelerated protocol (with “odd” suffix); resulting in SEMBodd, MESBdn, MEMBdn and MEMBodd. For each participant and protocol, we created a contrast to identify regions associated with semantic processing (semantic > control). Additionally for the decoding analysis, we modelled each block using separate regressors (12 semantic and 12 control) to obtain beta images per block.

The following additional parameters were applied to all models: six motion parameters as regressors of no interest; micro-time resolution set to number of slices and micro-time onset to number of slices/2; highpass filter of 128 s; AR(1) model to account for serial correlations; and we turned off SPMs automated mask threshold and supplied a brain mask extracted from individual T1.

#### 2.4.3. 2^nd^ level (between-participant) analysis

##### 2.4.3.1. Region of interest (ROI) analysis

###### 2.4.3.1.1. Univariate analysis

We had an a priori hypothesis that the semantic network would be activated during the semantic > control contrast. We took regions of interest from a large-scale spin-echo fMRI study that incorporates multi-model (visual and auditory) language tasks (Humphreys et al., 2015); however, given the large number of ROIs, we only inspected ones that had any overlap with the overall main effect in the present study (an F-test across all protocols). We used a custom MATALB script to extract ROI information built on SPM tools *(‘roi_extract.m’* from https://github.com/MRC-CBU/riksneurotools/Util). For each ROI, we extracted two metrics: 1) activation magnitude, i.e., the difference in GLM parameter estimates (“betas”) for semantic minus control blocks, and 2) activation precision, i.e., the t-statistic for that difference, i.e., the activation magnitude scaled by an estimate of the uncertainty of that difference, which also depends on the noise in the data. Then to determine the reliability of these two metrics, we performed planned t-tests across participants to compare protocol choices, i.e.: ME > SE (main effect of echo, i.e. [MEMB+MESB] - [SEMB+SESB]), MB > SB (main effect of band, i.e., [MEMB+SEMB] - [MESB+SESB]), the interaction between echo and band (i.e., [MEMB-SEMB] - [SEMB-SESB]), MEdn > ME (effect of denoising, i.e. [MEdnMB+MEdnSB] - [MEMB+MESB]) and MBodd > SB (effect of sampling rate, i.e. [MEMBodd + SEMBodd] - [MESB+SESB]).

We focused on two univariate effects: the effect of MR protocol on 1) activation magnitude (BOLD signal change) and 2) activation precision (reliability of BOLD change, analogous to CNR). The former was tested by comparing across MRI protocols the difference between the 1^st^-level betas for Semantic > Control conditions for each participant; the latter was tested by comparing the 1^st^-level T-statistic for this contrast instead (i.e., activation magnitude normalised by estimate of residual error across scans). A priori, one would expect ME to improve activation magnitude in regions prone to susceptibility artefacts by recovering signal drop-out, while one would expect MB to improve activation precision by having more data (effective degrees of freedom, dfs) and less aliasing of low-frequency fMRI noise (with little difference in activation magnitude). Downsampling the MB data (by dropping every even volume) should attenuate the MB advantage in activation precision due to higher dfs, but a residual advantage might remain owing to reduced noise aliasing. Likewise, reducing noise by removing using TE-dependent ICs from ME data should improve activation precision for ME protocols (but not affect activation magnitude, unless signal is removed by mistake).

###### 2.4.3.1.2. Decoding analysis

We also tested whether the protocols affected multivariate effects, namely decoding of condition using multivoxel pattern analysis (MVPA). For this, we extracted beta values within each ROI for each task block, which resulted in a 24 (12 semantic and 12 control blocks) x 280 (voxels per ROI) matrix. From this, we estimated the dissimilarity (using a cosine measure) between the patterns for every pair of blocks, and then averaged those according to whether they were within the same condition or from different conditions. We then subtracted the mean between-condition dissimilarity from the mean within-condition dissimilarity (which should be positive if decoding is possible), and used a non-parametric t-test to test whether this decoding metric differed for the contrasts across MR protocols described in the univariate section above. Decoding should be sensitive to factors that affect activation magnitude and activation precision (at individual voxels), and is potentially more sensitive than univariate analyses (Davis et al., 2014).

##### 2.4.3.2. Whole brain analysis

In order to determine if any effects were observed outside the a priori semantic network ROIs, we calculated whole brain effects using a 2x2 ANOVA to test for main effects of echoes and band. We used a custom MATLAB script built on SPM tools *(‘batch_spm_anova.m’* from https://github.com/MRC-CBU/riksneurotools/GLM) and extracted the following group level contrasts for semantics: 1) ME>SE; 2) SE>ME; 3) MB>SB; 4) SB>MB; and 5) the overall main effect of all protocols together. We applied a voxel-height threshold of p<0.001 uncorrected to define clusters, and then used a cluster-wise Family-Wise-Error corrected p<0.05 for statistical inference.

##### 2.4.3.3. Slice leakage analysis

To evaluate the potential impact of slice-leakage artefacts on this dataset, candidate artefact locations (artefact voxels) were identified for the MEMB and SEMB protocols, using a modified version of the MATLAB scripts developed for McNabb et al. 2016 (https://github.com/DrMichaelLindner/MAP4SL) (implemented in MATLAB 2020b). We conducted two types of analyses: 1) comparing univariate t-values; and 2) comparing multivariate decoding.

For the univariate analysis, we created GLMs on the pre-processed native EPI data (i.e., no normalisation or smoothing) and identified the voxel corresponding to the maximum t-value for each participant, separately for the SEMB and the MEMB data (seed voxels). For the SEMB protocol, GRAPPA was not used for in-plane acceleration and therefore only one alias location was expected per slice due to the MB protocol, with a phase-shift of FOV/2 (Figure 2 left). For the MEMB protocol, there were two alias locations per slice due to multi-banding and in-plane acceleration (Figure 2 right). An ROI of 3x3x3 voxels was then defined around each seed and corresponding artefact locations from which the mean t-value calculated for each ROI. The seed ROIs are labelled ‘A’ and the candidate artefact location due to multi-band is labelled ‘B’. ‘Ag’ represents the expected artefact location for ROI ‘A‘ due to GRAPPA, and ‘Bg’ is the equivalent artefact location for ROI ‘B’.

**Figure 2.**
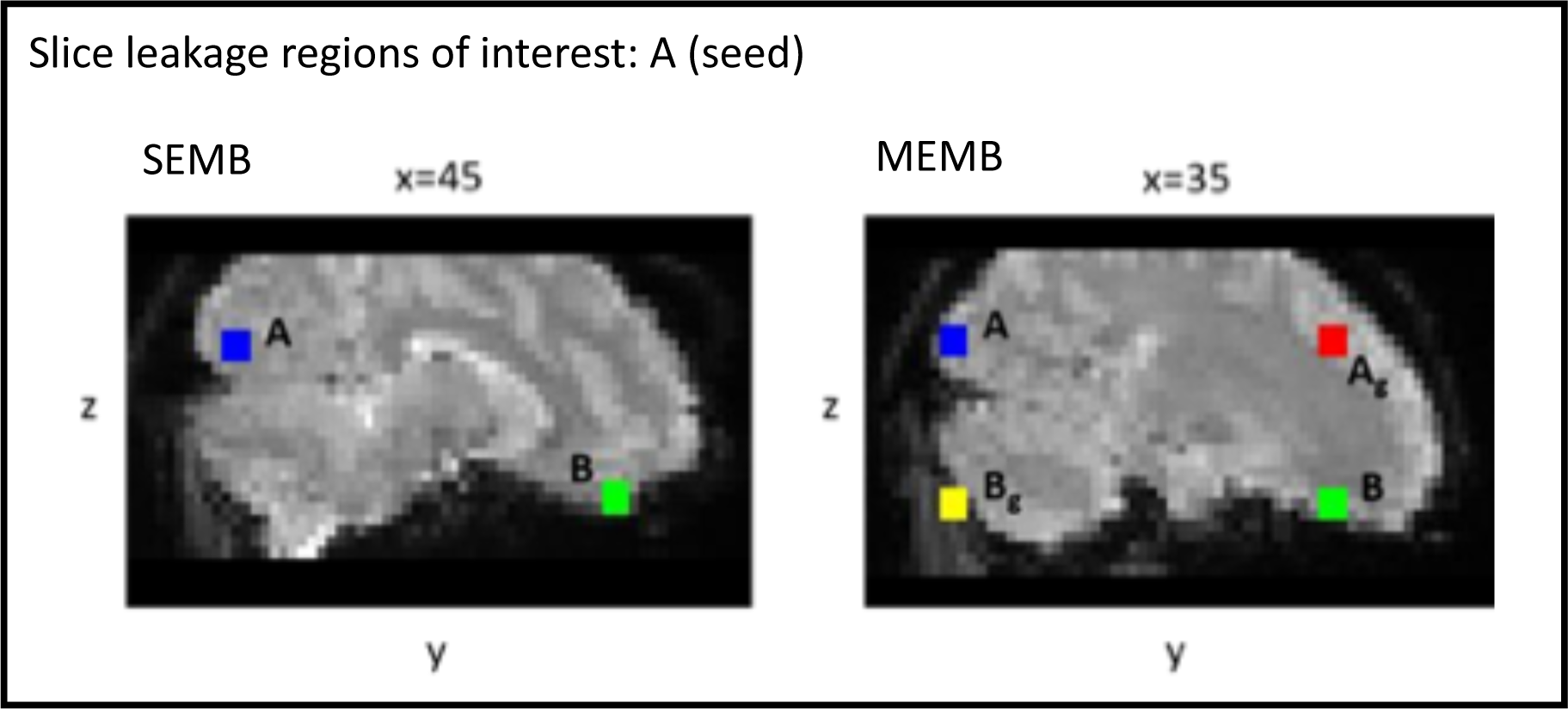
Example location of seed and artefact ROIs. Seed ROI [A, blue] and expected artefact locations for an individual participant. ROI B [green] represents the artefact location due to the multi-band protocol (phase shift of FOV/2), while ROI Ag [red] and Bg [yellow] represent the artefact locations due to in-plane acceleration (GRAPPA factor of 2). ROIs are shown for single echo and multi echo data.

For the multivariate analysis, we create GLMs on the EPI data in MNI space (but no smoothing), in order to have a greater number of voxels per ROI. The left vATL was used as the seed as this area showed successful decoding and is a key area of interest for the paradigm (see results in Table 2). Artefact locations were created in the same way as described above for the univariate analysis (i.e., in native EPI space) but all final ROIs were projected to MNI space and fixed as a sphere with 8mm radius.

**Table 2.** Showing t-test statistical comparisons between different planned contrasts using semantic network regions of interest (ROI) for activation magnitude (contrast of betas), activation precision (statistical T-values) and decoding (cosine dissimilarity). [*p<0.05, **p<0.05 corrected for 20 tests (5 ROIs x 4 comparisons)]. Abbreviations: inferior frontal pole (IFP), left ventral anterior temporal lobe (lvATL), left inferior frontal gyrus pars triangularis (lIFGtri), left posterior middle temporal gyrus (lpMTG), right inferior temporal gyrus (rITG), multi-echo (ME), single-echo (SE), multi-echo ICA denoised (MEdn), multi-band odd volumes (MBodd)

| Metric | Contrast | IFP | lvATL | lIFGtri | lpMTG | rITG |
| --- | --- | --- | --- | --- | --- | --- |
| Activation magnitude | ME>SE | 0.797 | **0.046\*** | 0.418 | 0.716 | **0.040\*** |
|  | MB>SB | 0.579 | **0.016\*** | 0.102 | 0.196 | 0.574 |
|  | MEdn>ME | 0.146 | 0.848 | 0.683 | 0.387 | 0.979 |
|  | MBodd>SB | 0.665 | 0.083 | 0.150 | 0.252 | 0.392 |
| Activation precision | ME>SE | 0.804 | 0.402 | 0.285 | 0.616 | 0.576 |
|  | MB>SB | **0.045\*** | **0.000\*\*** | **0.002\*** | **0.002\*** | **0.001\*\*** |
|  | MEdn>ME | 0.076 | **0.000\*\*** | **0.001\*\*** | 0.070 | 0.090 |
|  | MBodd>SB | 0.344 | **0.001\*\*** | 0.177 | 0.280 | 0.164 |
| Decoding | ME>SE | 0.889 | 0.581 | 0.162 | 0.143 | 0.697 |
|  | MB>SB | 0.310 | **0.001\*\*** | **0.004\*** | **0.006\*** | 0.509 |
|  | MEdn>ME | 0.865 | **0.001\*\*** | **0.024\*** | 0.245 | **0.047\*** |
|  | MBodd>SB | 0.281 | 0.119 | 0.051 | 0.138 | 0.970 |

To assess whether false positive activations/decoding due to slice-leakage could be observed in the multi-band data, the corresponding single-band data (SESB and MESB, respectively) were used as controls since no aliasing artefacts are expected. For this purpose, we generated seed and artefact ROIs (as defined for SEMB) in the SESB data and extracted t-values/dissimilarity. Similarly, the seed and artefact ROIs generated for the MEMB data were used to extract t-values/dissimilarity from the MESB data

## 3. Results

### 3.1. Behavioural data

We identified four participants who had poor behavioural performance (2SD away from group mean, where mean accuracy of all conditions < 73% across all runs), and therefore removed them from further analysis. We also excluded an additional participant who had poor EPI coverage of the temporal lobes. The final sample of 16 participants had good and comparable performance (Supplementary Materials 1). The results for the ANOVA’s showed that behavioural performance was consistent across all protocols, even though the semantic task was less accurate than the control task. The ANOVA for accuracy revealed a main effect of condition (F(1,15)=131.086, p<0.001) (78.46 [SD=6.37] vs 94.03 [SD=7.13]% for semantic and control trials, respectively). Both conditions were significantly above chance (p’s<0.001). No other effects or interactions were observed for accuracy. We did observe a 3-way interaction for RT (F(1,15)=6.566, p=0.022); however, follow-on ANOVAs broken down by one of the factors showed no significant effects, so this 3-way interaction is difficult to interpret further and no other effects were significant.

### 3.2. ROI results

The ROIs from Humphreys et al. (2015) that overlapped with significant clusters for the contrast of Semantic vs Control conditions, averaged across MR protocols, are shown in Figure 3. These included left ventral anterior temporal lobe (vATL), right inferior temporal gyrus (ITG), left inferior frontal gyrus (IFG), left posterior middle temporal gyrus (pMTG) and left frontal pole (FP).

**Figure 3.**
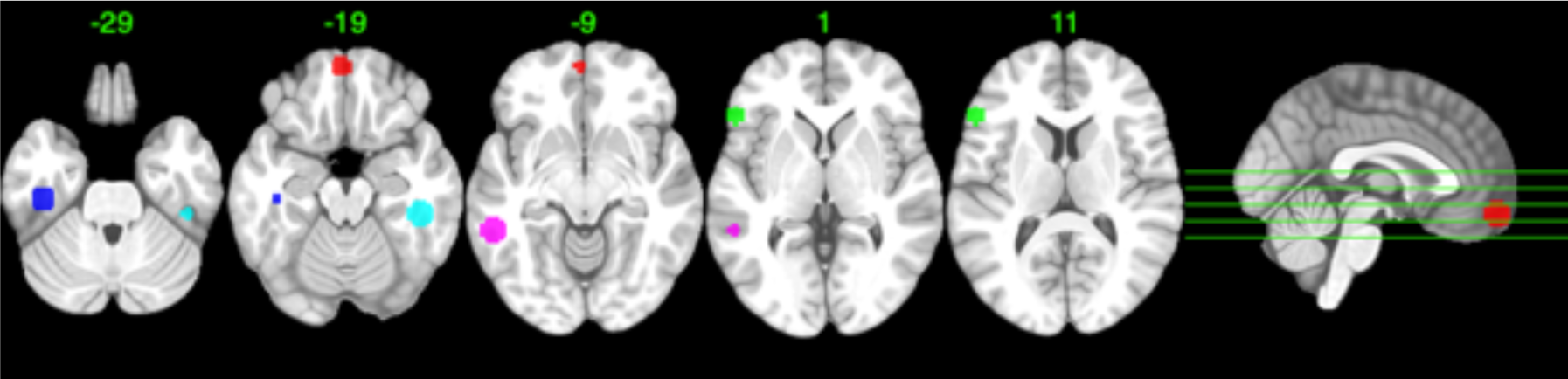
Semantic network regions of interest (ROI; 8mm spheres) defined using Humphreys et al. (2015) and selecting ones that overlap with overall main effect of semantics in the current study. Inferior frontal pole (red), left ventral anterior temporal lobe (blue), left inferior frontal gyrus pars triangularis (green), left posterior middle temporal gyrus (purple), right inferior temporal gyrus (cyan)

The planned contrasts of interest and resultant statistical p-values are shown in Table 2 (note see Supplementary Materials 1 for an extended table of comparisons, including reverse contrasts). For activation magnitude (contrasts on the mean across participants of the difference between S and C betas), ME was marginally higher than SE in left vATL and right ITG. This would be expected if these regions suffer from susceptibility artefacts, causing greater “drop-out” of signal for the SE with non-optimal TE for those regions. For MB, we also found a significant benefit over SB in the left vATL. The ICA-denoising (MEdn) did not improve on the basic ME data. The interaction between echo and band was not significant in any ROI. We show results across a wider ROI network and the reverse contrast in Supplementary Materials 1; note we found one effect in the opposite direction, where ME > MEdn in right ITG (p=0.021) (see Supplementary Materials 1).

For activation precision (comparisons across participants of their T-statistics), there were advantages for MB vs SB for all 5 ROIs, but no differences for ME vs SE. The former main effect of MB is expected from the greater sampling rate (i.e, shorter TR), resulting in more effective degrees of freedom in the data and less aliasing of low-frequency noise. As expected, when removing one half of the (even) volumes (MBodd), this advantage disappeared for most regions, though a MB advantage remained in lvATL. Though there was no evidence for a basic advantage of ME versus SE, once ICA-denoising was applied (enabled by having more than one TE), the activation precision increased (for MEdn versus ME) in two of the ROIs, suggesting that noise sources are successfully being removed. We did not find any significant results for reverse comparisons for these ROIs (see Supplementary Materials 1).

Finally, the decoding results were very similar to the activation precision results, with no evidence for a benefit for ME over SE protocols, but superior decoding for MB over SB protocols in the left vATL, left IFGtri, and left pMTG. Similarly, ICA-denoising improved data decoding effects in the left vATL, left IFGtri, and right ITG. The matched sampling MB data marginally outperformed the SB protocols in the left IFGtri only. As above, the reverse comparisons showed fewer differences, with an SB advantage observed over the reduced MB time-series only in the rITG (p=0.030).

### 3.3. Whole-brain results

For activation magnitude, Figure 5a shows whole-brain results for the semantic>control contrast, averaged across all protocols (the results for individual protocols are provided in Supplementary Material 3). Activations are largely bilateral, and include the core semantic regions in the temporal and frontal cortices. Results for the main effects of ME and of MB in the 2x2 ANOVA are shown in Figure 5b and 5c, with peak information is summarised in Supplementary Materials 4. The main effect of Echo identified three clusters with greater activation magnitude for ME than SE: 1) left inferior temporal/fusiform cortex, 2) left anterior cingulate gyrus and 3) right frontal pole. By contrast, clusters in bilateral medial temporal fusiform, parahippocampal and hippocampal regions showed the opposite effect of greater magnitude for SE than ME. Finally, the main effect of band revealed greater activation magnitude for MB than SB in similar bilateral medial temporal structures such as temporal fusiform cortex, with no significant clusters for the reverse contrast. No voxels survived correction for the ME-by-MB interaction.

**Figure 5.**
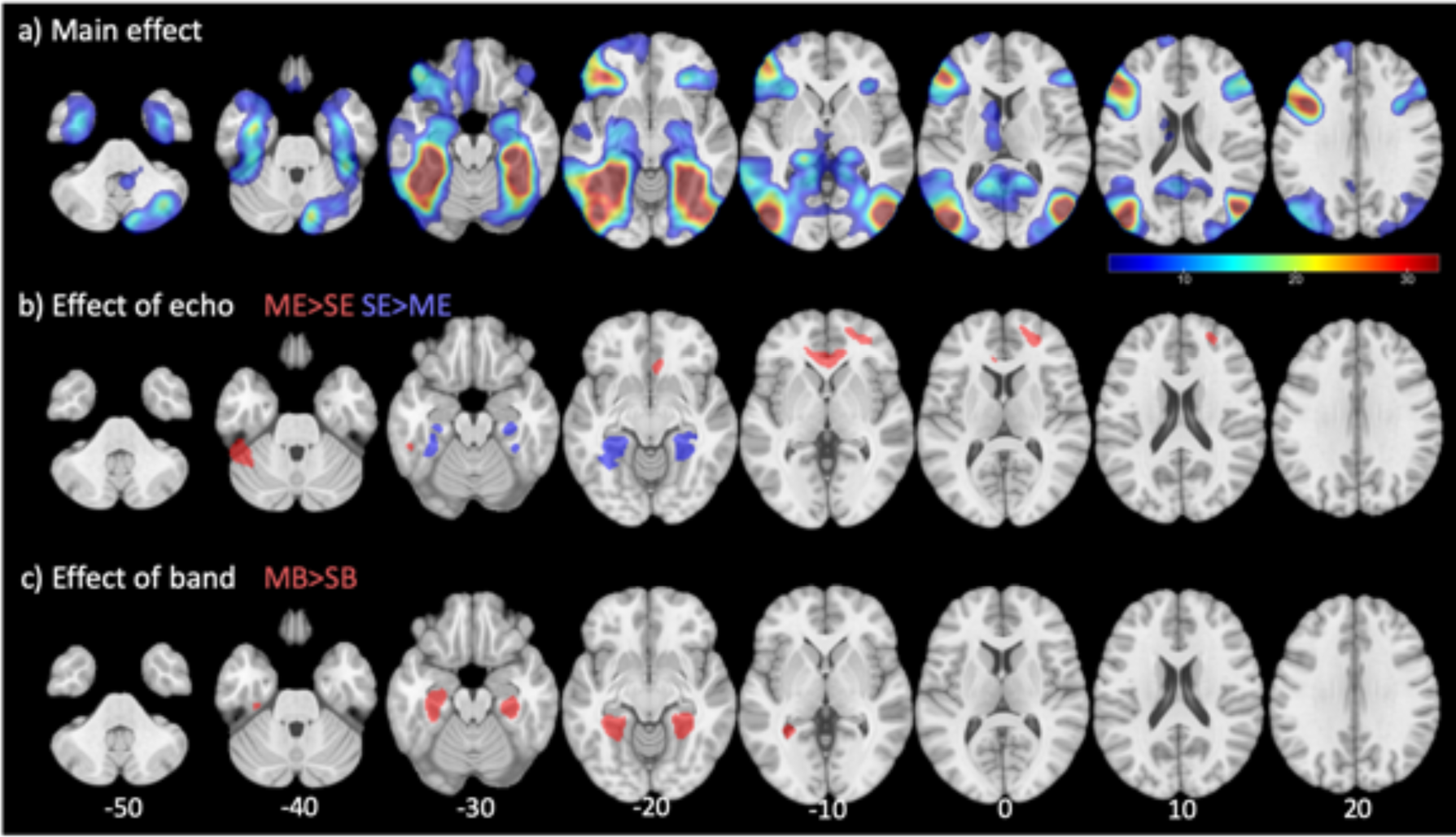
Whole brain analysis comparing activation magnitude (contrast betas) for Semantic > Control in a 2x2 ANOVA manipulating echo and band fMRI protocols. a) average effect across all protocols [t-value 3.28-32.8]; b) directed effects of echo (ME>SE [red] and SE>ME [blue]); and c) directed effects of band (MB>SB [red]; SB>MB not significant)

The results using activation precision (Figure 6) were largely similar to those for activation magnitude. The main noticeable difference was that MB was significantly better than SB in almost all parts of the semantic network, including extending into the ventral anterior temporal fusiform cortex.

**Figure 6.**
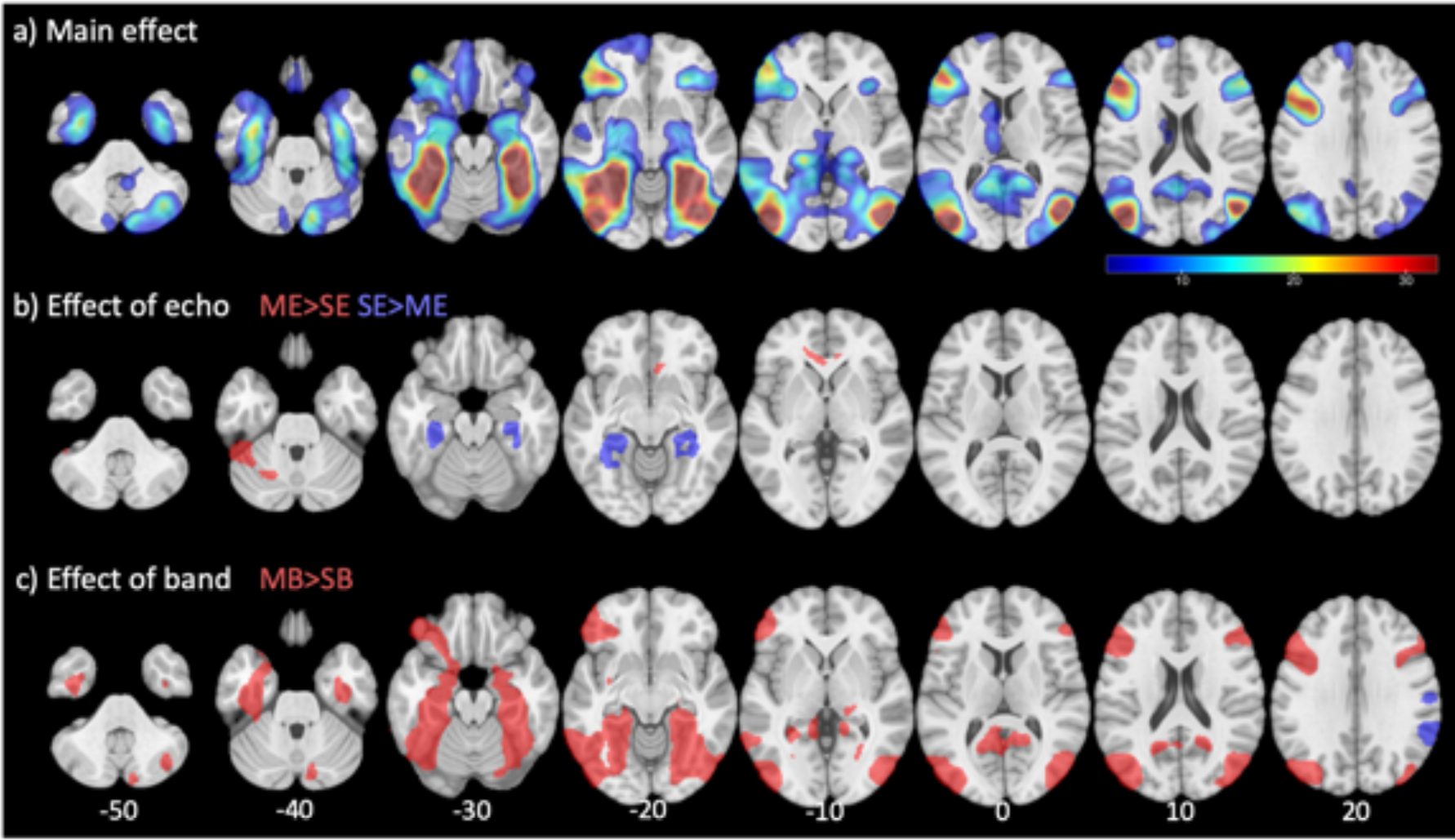
Whole brain analysis comparing activation precision fit (statistical t-values) during semantic activation (semantic>control) in a 2x2 ANOVA manipulating echo and band fMRI protocols. a) overall main effect of all protocols [t-value 3.28-32.8]; b) directed effects of echo (ME>SE [red] and SE>ME [blue]); and c) directed effects of band (MB>SB [red] and SB>MB [blue])

We also ran a 2x2 ANOVA on the multi-echo data only to test the effect of ICA-denoising and whether this interacted with multi-band. We observed only one cluster surviving correction for activation magnitude, where the main effect of band (MB>SB) identified left posterior fusiform cortex clusters extending up into the right calcarine sulcus (Supplementary Materials 5a). Differences in activation precision were similar to those shown in Figure 6, where the MB>SB contrast showed clusters across the semantic network (Supplementary Materials 5b). Overall, this analysis replicated results showing that MB offers improvements over SB across the entire semantic network; in contrast ICA-denoising did not show specific improvements but was not significantly worse than the t2star dataset (i.e., with no ICA).

### 3.4. Slice leakage results

Supplementary Materials 6(i) shows the mean t-value per participant for seed and artefacts ROIs for the single-echo protocols, and Supplementary Materials 6(ii) shows the data for the multi-echo protocols. For both univariate and multivariate analysis, we used paired t-tests to compare the mean t-values/dissimilarity between corresponding ROIs for single-echo protocols (t_A,SEMB_ – t_A,SESB_ and t_B,SEMB_ – t_B,SESB_), and for multi-echo protocols (t_A,MEMB_ – t_A,MESB_, t_B,MEMB_ – t_B,MESB_, t_Ag,MEMB_ – t_Ag,MESB_ and t_Bg,MEMB_ – t_Bg,MESB_).

For the univariate analysis, the difference for the seed ROI (A) is significant between SEMB and SESB (t=5.817, df=20, p<0.001) and also between MEMB and MESB (t=4.405, df=20, p<0.001), reiterating the previous results where MB results in higher t-values than SB protocols. For the artefact locations, no comparison was significant, with the exception of ROI Bg for the multi-echo data, where the mean t-values were found to be significantly greater for MESB when compared to MEMB (t=-2.096, df=20, p=0.048). However, the directionality is opposite to what we might expect for slice-leakage artefacts, and the comparison does not survive correction for multiple comparisons.

For the multivariate analysis, the difference for the seed ROI (A) is significant between SEMB and SESB (t=3.295, df=20, p<0.005) but not between MEMB and MESB (t=0.2488, df=20, p<0.8069). For the artefact locations, no comparison was significant. Therefore, overall the results suggest no evidence of slice-leakage effects for univariate or multivariate analysis in the current dataset.

## 4. Discussion

Gradient-echo BOLD sensitivity varies across the brain. In particular, the presence of air cavities near the front and lateral sides of the head result in magnetic susceptibility artefacts (signal loss and distortions). Many methods have tried to improve signal detection in such areas while retaining sufficient sensitivity to the rest of the brain, and this study provides the first systematic comparison of a typical imaging protocol with multi-echo and multi-band modifications. When comparing the precision with which activations were detected during a semantic task (i.e, average T-statistics), we found that multi-band protocols were generally beneficial, with no evidence of signal leakage artefacts, at least for a multi-band factor of 2. This increased precision is to be expected given the higher number of volumes and hence additional degrees of freedom with which to estimate activation. Also as expected, when comparing the magnitude of activations, multi-echo protocols increased activations in regions prone to susceptibility artefacts (specifically the anterior temporal lobes, ATLs), and the addition of ICA-denoising (enabled by measuring multiple echoes) further improved the precision of those activations. Both multi-banding and ICA-denoising of ME data also tended to improve multi-voxel decoding of experimental conditions. However, multi-echo protocols reduced activation magnitude in more central regions, such as the medial temporal lobes (MTLs), presumably due to the higher in-plane acceleration (GRAPPA) required to record multiple echoes.

### 4.1 Advantages and disadvantages of multi-echo protocols

In the current study, we probed semantic cognition with task-based fMRI. There is substantial evidence supporting the hub-and-spoke theory of semantic representation, where the ATLs serve to integrate information from multiple modality specific spokes (Lambon Ralph et al., 2017). Detecting fMRI activation for semantics in the ventral and lateral ATLs has been challenging, due to a number of factors (reviewed in Visser, Jefferies, & Lambon Ralph, 2010) and chief among them is likely to be signal loss with conventional TEs ∼30-35ms. As noted in the Introduction, modified protocols have been successful at detecting activity using spin-echo fMRI (Binney et al., 2010; Humphreys et al., 2015; Visser, Embleton, et al., 2010) and dual gradient-echo fMRI (Halai et al., 2015, 2014). In the current study, all protocols were able to detect robust activations in bilateral ventral ATL as well as inferior frontal regions. Nonetheless, we demonstrated that certain modifications can lead to further improvements.

A priori, we expected multi-echo to improve activation magnitude in regions prone to susceptibility artefacts by recovering signal drop-out, compared to single-echo protocols. In our ROI analysis, we observed such a benefit in the left ATL and right ITG for activation magnitude. Interestingly, this increased signal was not accompanied by higher T-statistics (activation precision) – nor better multivoxel decoding – in these ROIs, suggesting that the ME protocol also increased noise (e.g, due to the in-plane acceleration required to reduce echo train length). Nonetheless, when leveraging the multiple echoes to improve ICA-denoising, these ROIs did now show increased activation precision and multivoxel decoding (see Supplementary Materials 2).

Signal recovery in areas near susceptible artefacts are expected with multi-echo protocols because the shorter TEs are better able to capture signal before it dephases. This is in-line with current language literature using similar modified protocols (i.e., Halai et al., 2014; Jung et al., 2017). When using a whole-brain search, we did see increased activation magnitude and precision in a few other clusters, most notably left posterior inferior temporal gyrus, which is also prone to susceptibility artefacts. Improvements were also seen in left anterior cingulate gyrus and right frontal pole, though the reason for this is less clear, given that these regions are not normally associated with susceptibility artefacts. Importantly, we also observed a significant reduction in both activation magnitude and activation precision in more central brain regions, such as the MTL. This is most likely due to lower tSNR in brain regions that are further from the receiver coils, which is exaggerated when using higher g-factors for in-plane acceleration (Kirilina et al., 2016). Although, Fazal et al., (2023) showed that multi-echo multi-band had reduced sensitivity in visual and anterior cingulate regions compared to multi-band only during a Stroop task.

Finally, we hypothesised that removing noise from the timeseries should improve activation precision and decoding (but not affect activation magnitude, unless signal is removed by mistake). MR physics dictates that there is a positive linear relationship between TE and BOLD signal amplitude, but not noise amplitude, and this fact is used by automated techniques that deploy ICA and machine learning to separate components likely to be signal from those likely to be noise, such that noise components can then be projected out of the data (DuPre et al., 2021; Kundu et al., 2013; Kundu, Inati, Evans, Luh, & Bandettini, 2012). Many studies have demonstrated the benefit of this “ME-ICA” approach for resting-state fMRI connectivity (e.g., Cohen, Yang, et al., 2021; Lombardo et al., 2016; Lynch et al., 2020); few however have demonstrated this benefit in task-based fMRI. Here we also failed to find any clusters in our whole-brain analysis that survived correction for the main effect of ICA-denoising (for ME protocols averaged across SB/MB; Supplementary Material 5). This suggests that effects of ICA-denoising are generally weak in this task. Nonetheless, we did find evidence that ICA-denoising improves task-related activation precision and decoding when using more sensitive tests within some of our a priori ROIs. These improvements presumably reflect a reduction in overall residual error (noise). Interestingly, one ROI (right ITG) actually showed a reduction in activation magnitude after ICA denoising (see Supplementary Material 2), suggesting that some signal of interest may have been removed by mistake. In sum, we propose that automated ICA denoising methods that use the TE-dependence of BOLD enabled by ME protocols can improve task-based statistics, but only modestly, and the risk of also removing signal should be kept in mind.

### 4.2 Advantages of multi-band data

For the multi-band modification, we expected improvements to activation precision owing to having more data and less aliasing of high-frequency fMRI noise, without any impact on activation magnitude. There is some evidence that multi-band protocols can impair sensitivity when combined with in-plane acceleration or reduced k-space sampling (e.g., Chen et al., 2015), but here we were careful to match these to the single-band protocols (apart from the flip-angle, which was reduced so as to maximise BOLD sensitivity for the corresponding TR). Multi-band modifications have mainly been promoted for improved estimation of resting-state connectivity (Smith, Beckmann, Andersson, Auerbach, Bijsterbosch, Douaud, Duff, Feinberg, Griffanti, Harms, Kelly, Laumann, Miller, Moeller, Petersen, Power, Salimi-Khorshidi, Snyder, Vu, Woolrich, Xu, Yacoub, Uğurbil, et al., 2013; Uğurbil et al., 2013); their benefit for task-based fMRI analysis has been less clear (Demetriou et al., 2018; Todd et al., 2017), since the reduced low-frequency noise is unlikely to be correlated (phase-locked) with the task regressors, and so can be removed by standard high-pass filtering. Nonetheless, the present multi-band modification led to consistent benefits in activation precision, but not activation magnitude, across all a priori ROIs (Table 2). It also improved multivoxel decoding in many of them. Consistent with the hypothesis that this advantage arises primarily from more volumes (Constable & Spencer, 2001; Miller, Bartsch, & Smith, 2015), these improvements were largely removed when we sub-sampled only odd-numbered volumes. The only exception to this pattern was lvATL, which showed increased activation precision for MB protocols. The same increase in activation magnitude with multi-banding was also seen in MTL in the whole-brain analysis. The reason for these increased signal magnitudes is unclear – it could reflect an interaction between MB and ME modifications, but tests of this interaction did not reach significance.

Another known issue with increasingly high multi-band acceleration is slice leakage artefacts (Todd et al., 2016). However, we found no evidence of signal deviations (for both univariate or multivariate analyses) in areas expected to exhibit leakage for either single or multi-echo protocols. This could be due to the fact that an acceleration factor of 2 was used in this study, where Todd and colleagues reported significant leakage with acceleration factors greater than 4.

Finally, it is interesting that we found no significant interactions between ME and MB modifications, in either the ROI or whole-brain analyses. Thus there was no evidence that ME and MB are synergistic; their effects appear to be additive.

## 5. Conclusions

We showed that modifications to a typical fMRI protocol can lead to benefits in activation magnitude, precision and decoding through multi-echo and/or multi-band methods. In general, we found that multi-band proved to be beneficial for activation precision (T-statistics) and multivoxel decoding across many ROIs, whereas the multi-echo was mainly beneficial in areas affected by susceptibility, and improved activation magnitude. We observed some loss in quality for multi-echo methods in more central parts of the brain. Nonetheless, the MEMB protocol used here is a promising default option for fMRI on most brain regions, particularly those that suffer from susceptibility artefacts, as well as offering the potential to apply advanced post-processing methods to take advantage of the increased temporal (or spatial) resolution of MB protocols and more principled ICA-denoising based on TE-dependence of BOLD signals.

## Data and code availability

Data will be made publicly available upon peer review and acceptance. Code is publicly available at https://github.com/AjayHalai/Sensitivity_of_MEMB.

## Author contributions

AH, RH, MMC designed and conceptualised the study. AH, PF collected and curated the data; AH, RH, MMC conducted the analysis; AH, MMC prepared figures/tables; AH wrote the manuscript draft; AH, PF, RH, MMC reviewed and edited final manuscript.

## Acknowledgements

We would like to thank the participants and MRC Cognition and Brain Sciences Unit radiographers. AH was supported by a Career Development Award from the MRC (MR/V031481/1). RH was supported by MRC programme grant SUAG/086 G116768. MMC was supported by MRC Unit grant SUAG/019 G116768. For the purpose of open access, the author has applied a Creative Commons Attribution (CC BY) licence to any Author Accepted Manuscript version arising from this submission.

## Supplementary Materials

**Supplementary Materials 1.**
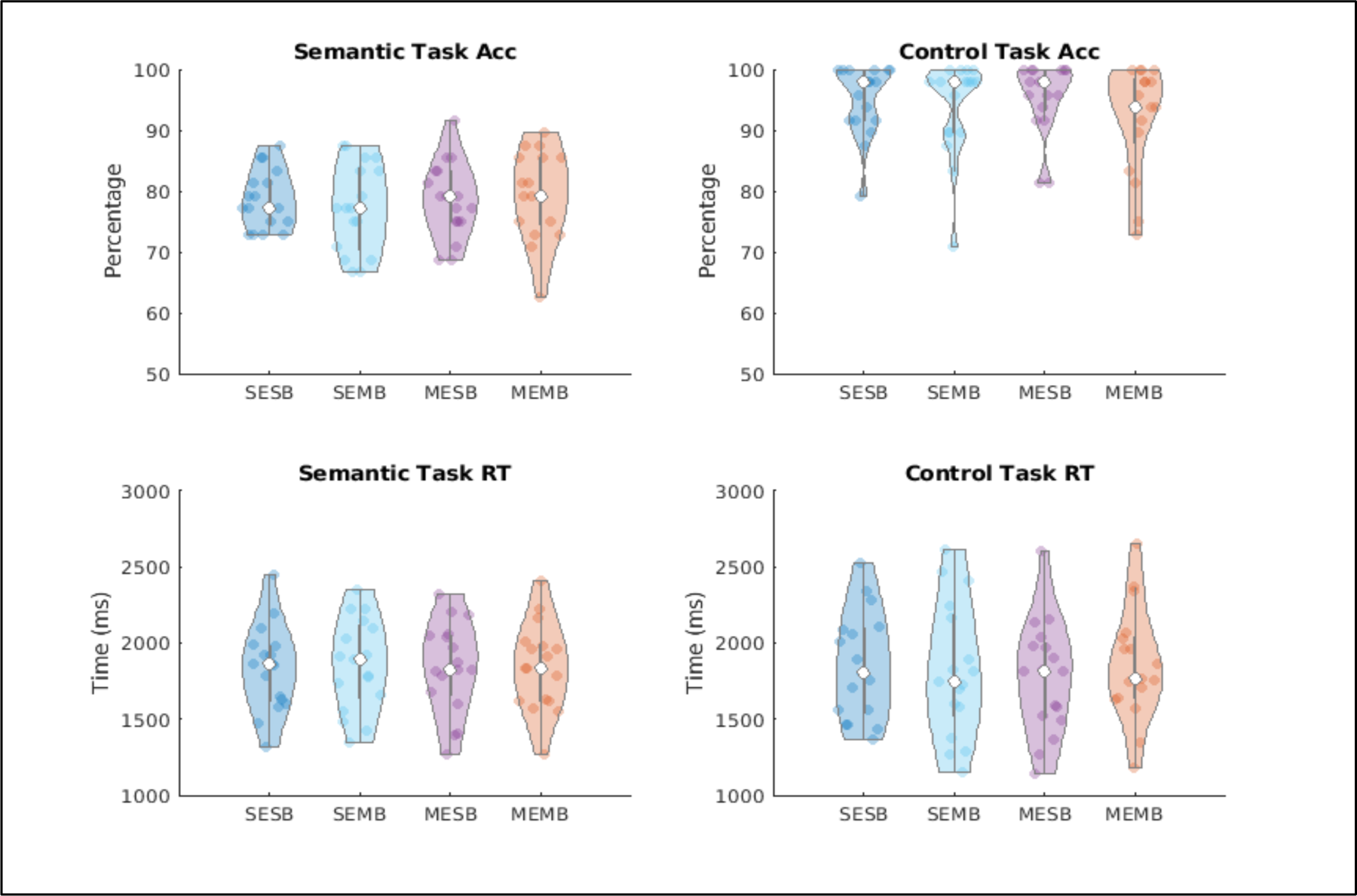
Accuracy and reaction time data for semantic and control trials. Overall, the figure shows consistent behavioural performance across protocols. The top row shows lower accuracy for semantic compared to control trials, whereas reaction time did not differ. Abbreviations: Single echo single band (SESB), single echo multi band (SEMB), multi echo single band (MESB), multi echo multi band (MEMB).

**Supplementary Materials 2.**
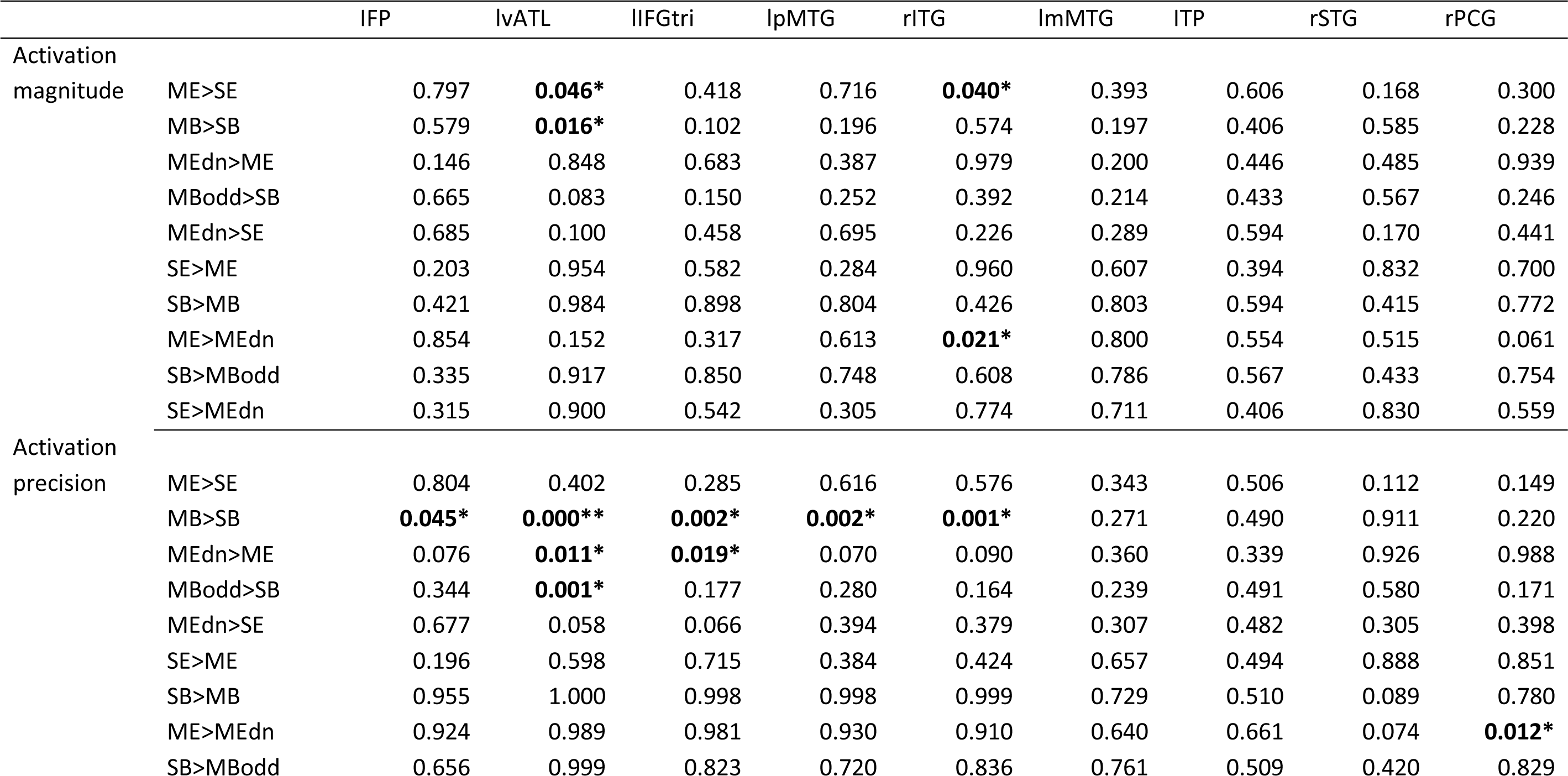
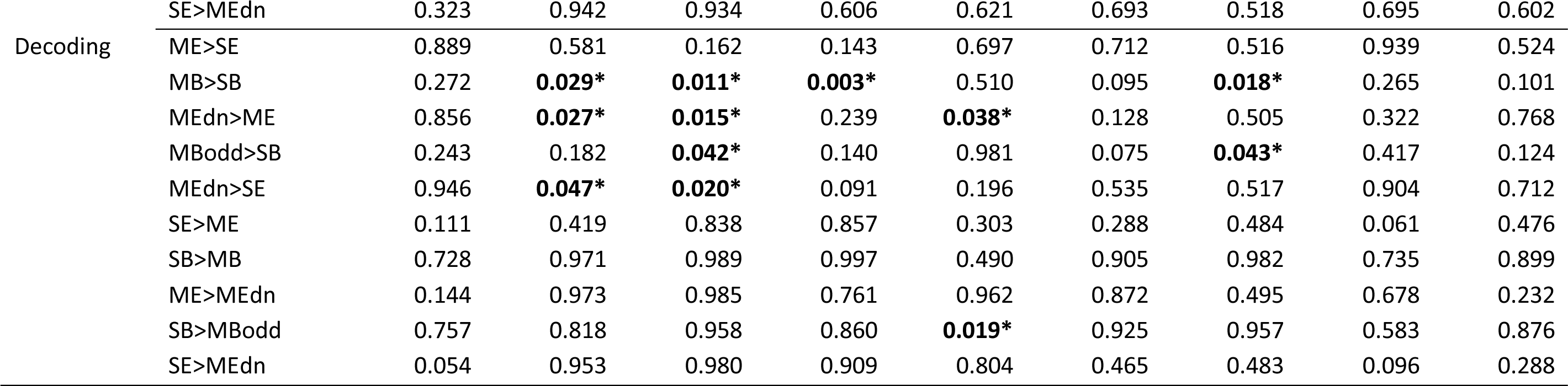
Table. Showing statistical comparisons across semantic network regions of interest (ROI) for activation magnitude (contrast of betas), activation precision (statistical T-values) and decoding (cosine dissimilarity). Note, columns here are ROIs from the full semantic network identified in Humphreys et al., 2015. [*p<0.05, **p<0.05 corrected].

**Supplementary Materials 3.**
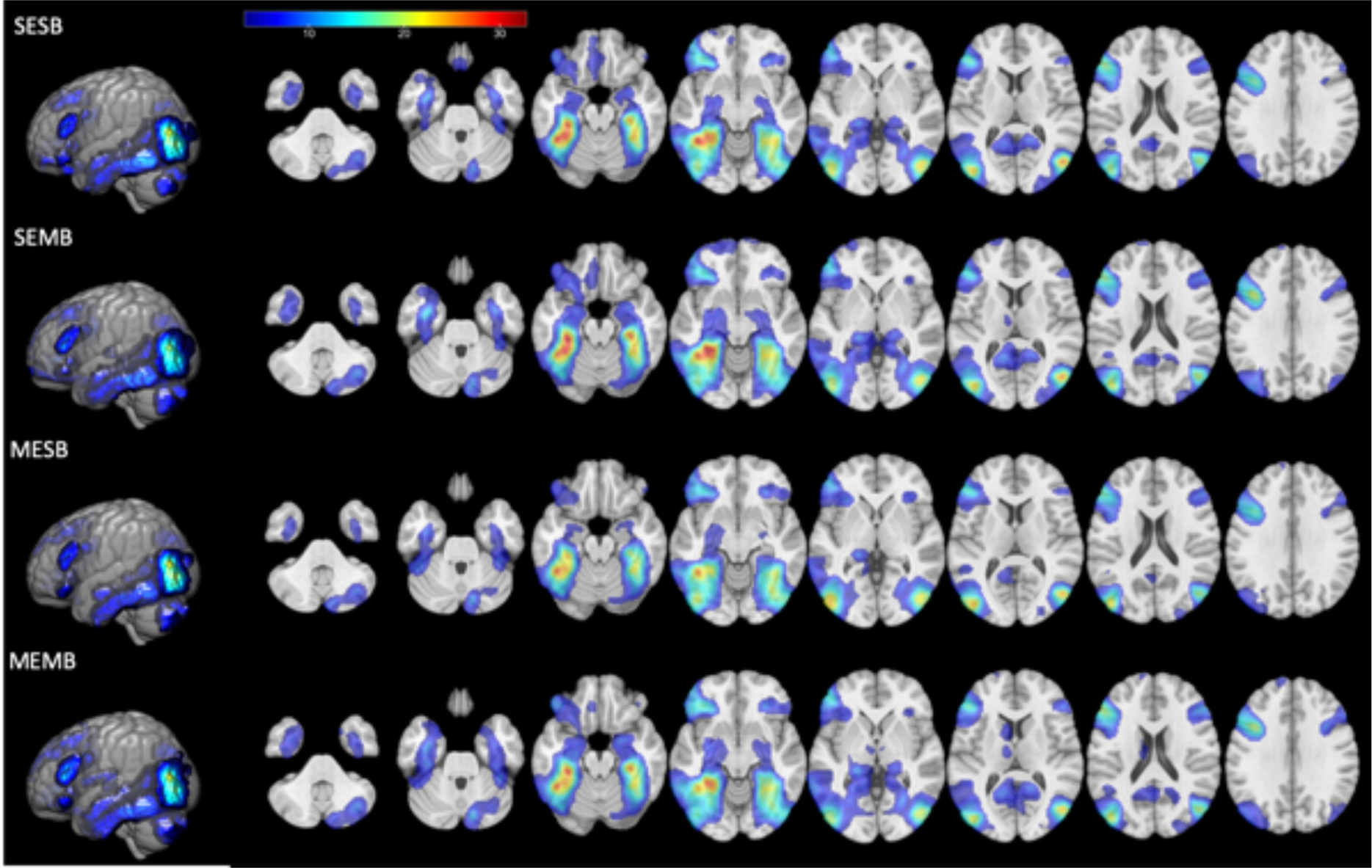
Whole brain results for the Semantic>Control contrast for each protocol [t-value 3.28-32.8], thresholded at p<0.001 voxel height FWE-cluster corrected p<0.05. Abbreviations: Single echo single band (SESB), single echo multi band (SEMB), multi echo single band (MESB), multi echo multi band (MEMB).

**Supplementary Materials 4.**
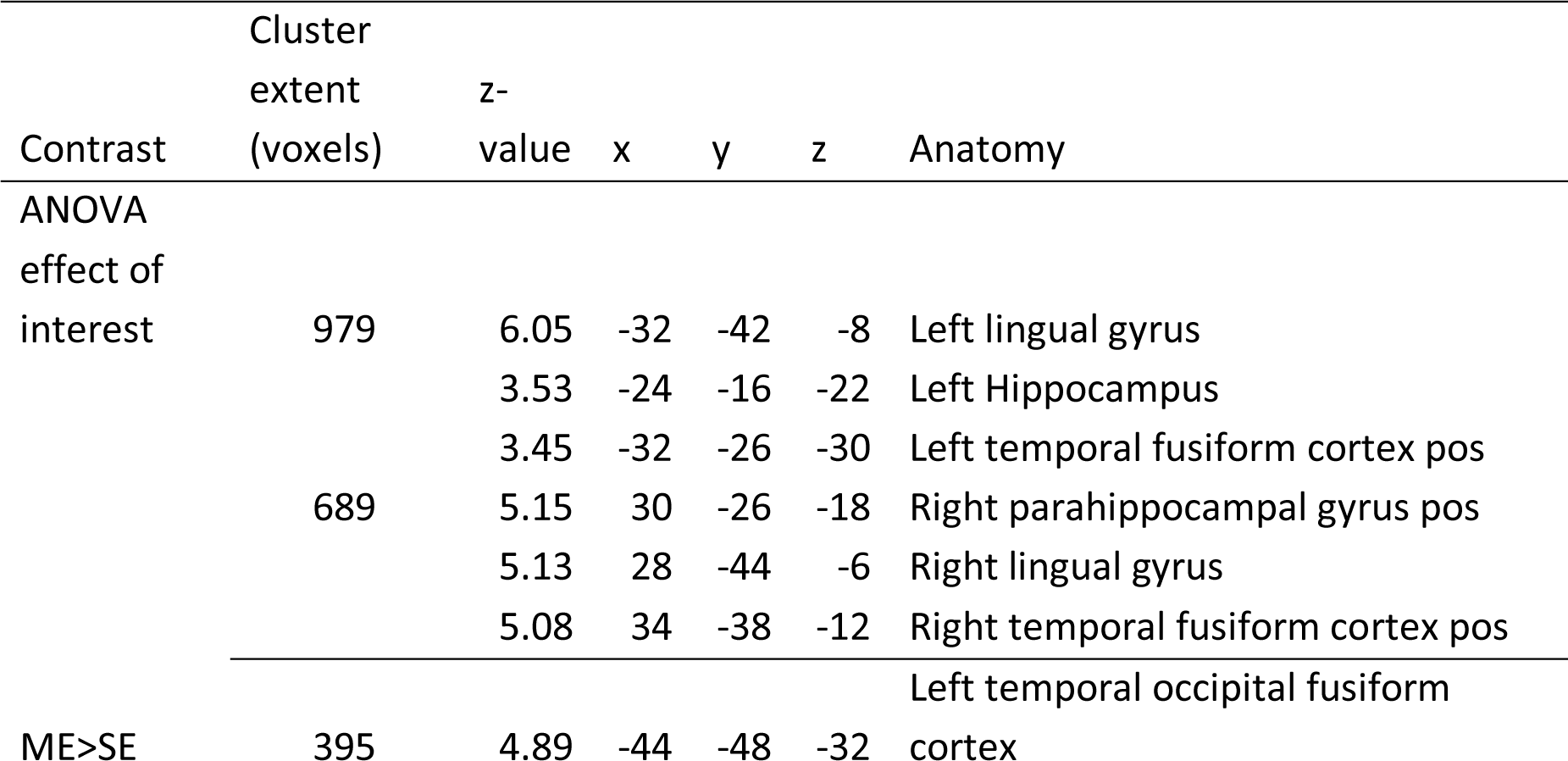
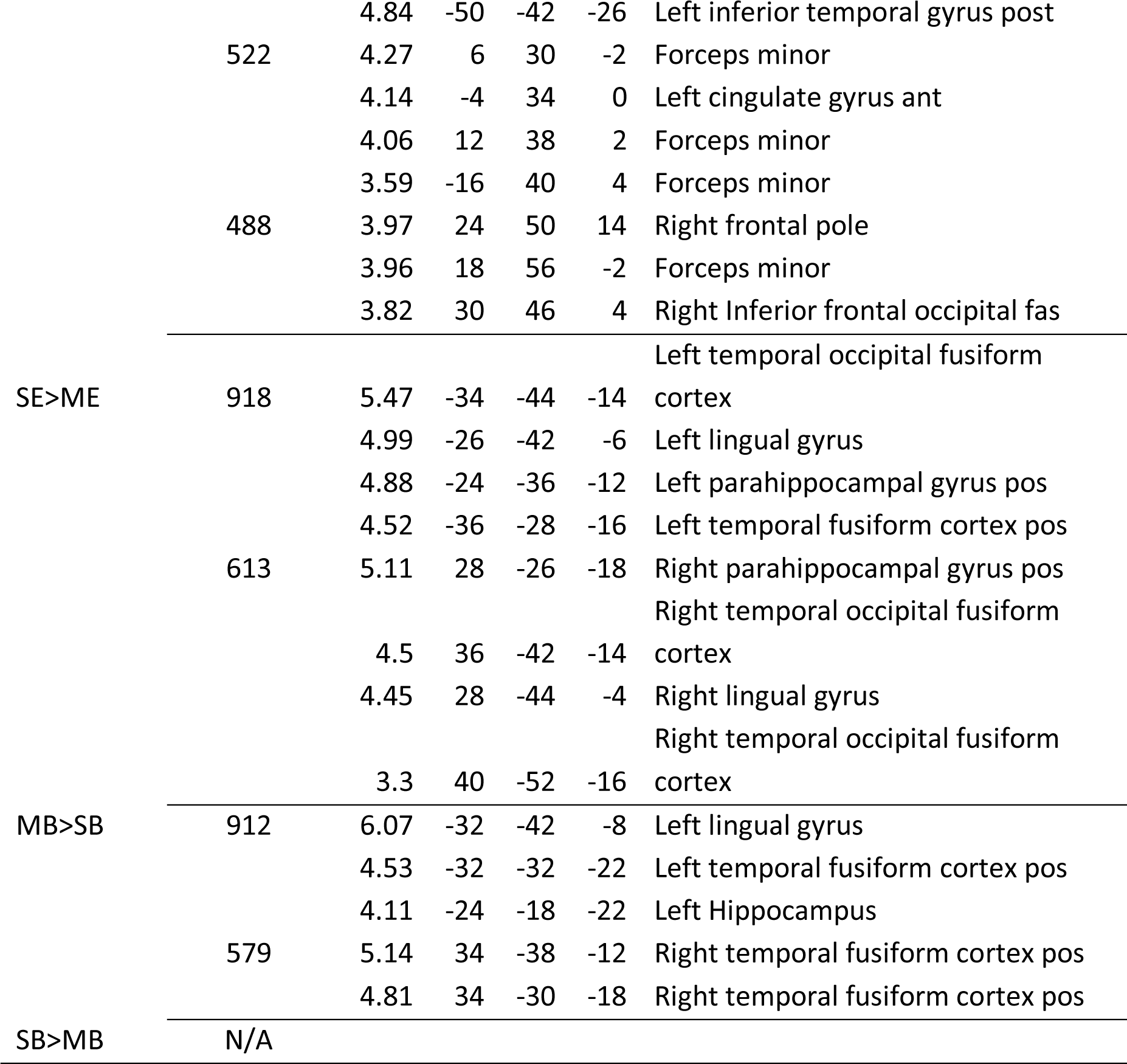
Table. Showing significant cluster and peak information for select contrasts of interest when comparing activation magnitude.

**Supplementary Materials 5.**
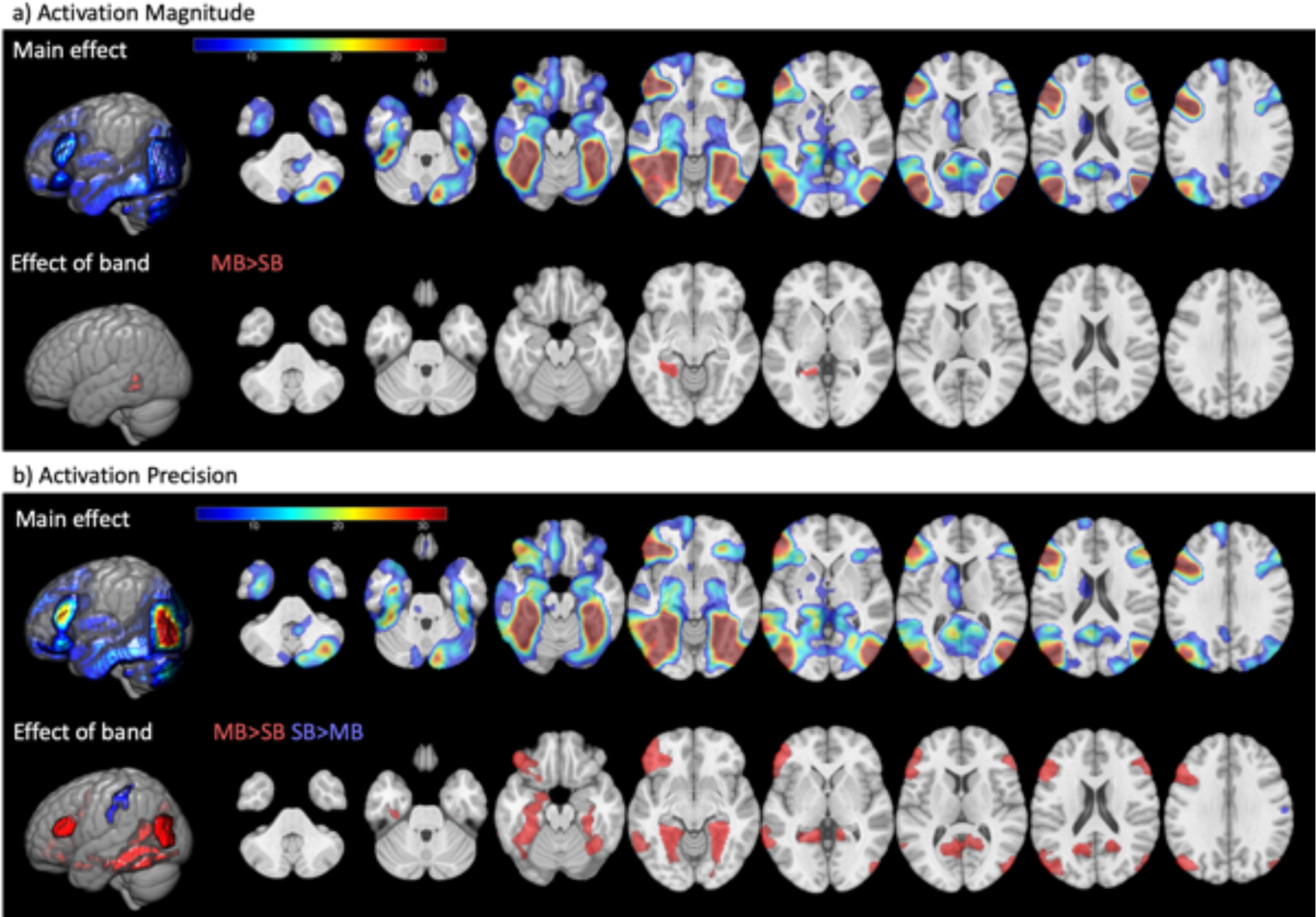
Whole brain results for the effect of ICA-denoising and band (2x2 ANOVA) on the multi-echo data. 4a) Results for activation magnitude (contrast betas) and 4b) for activation precision (statistical t-values) for semantic>control. Both sections show the average effect across all protocols [t-value 3.28-32.8] and directed effects of band (MB>SB [red] and SB>MB [blue]). Note that no clusters survived correction for the main effect of ICA-denoising. Abbreviations: Multi band (MB), single band (SB).

**Supplementary Materials 6.**
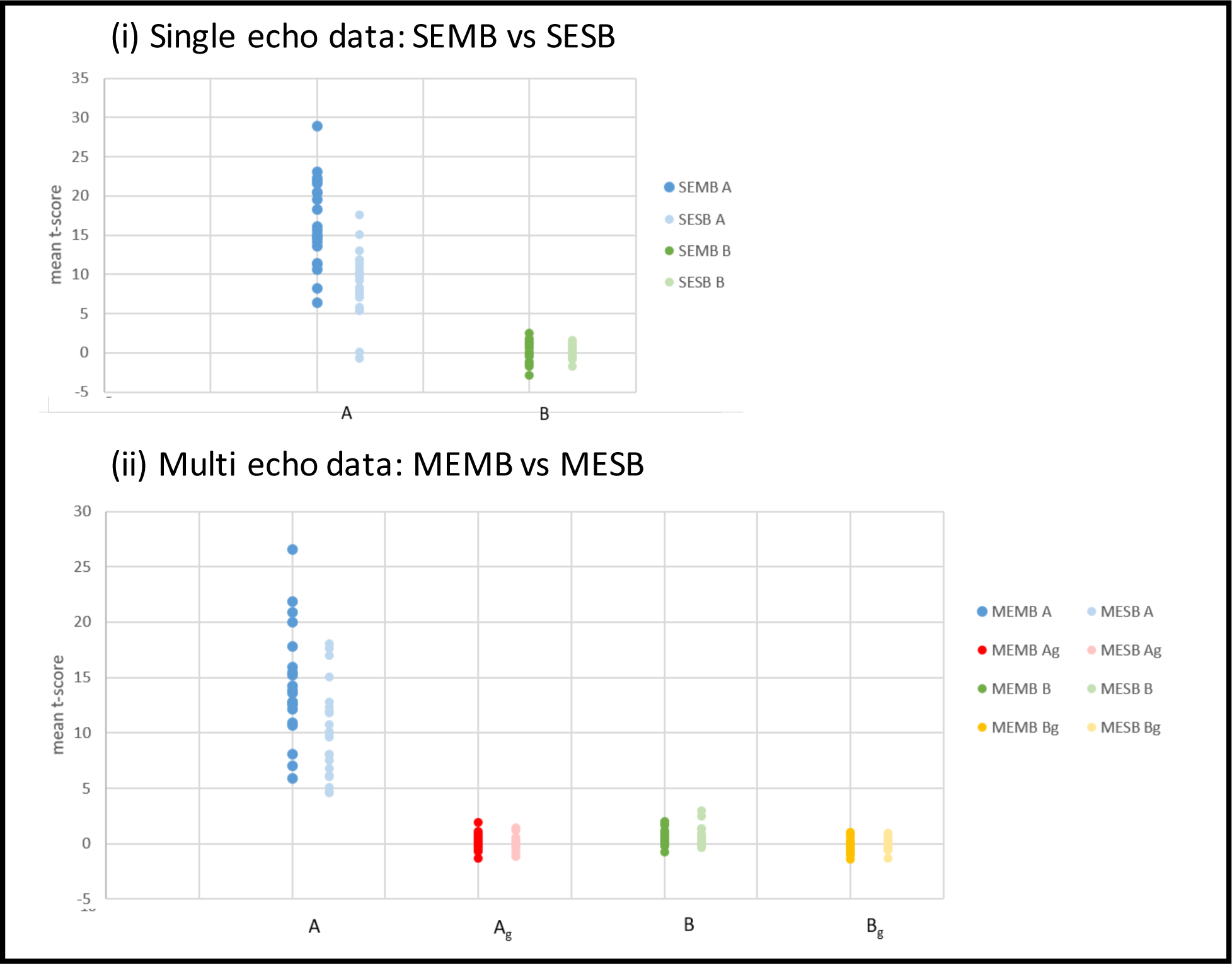
Mean t-values obtained for each subject, per seed and artefact ROI. Mean t-values for (i) SEMB and SESB, and (ii) MEMB and MESB data. Abbreviations: Single echo single band (SESB), single echo multi band (SEMB), multi echo single band (MESB), multi echo multi band (MEMB).

